# Morphological constraints on cerebellar granule cell combinatorial diversity

**DOI:** 10.1101/212324

**Authors:** Jesse I. Gilmer, Abigail L. Person

## Abstract

Combinatorial expansion by the cerebellar granule cell layer (GCL) is fundamental to theories of cerebellar contributions to motor control and learning. Granule cells sample approximately four mossy fiber inputs and are thought to form a combinatorial code useful for pattern separation and learning. We constructed a spatially realistic model of the cerebellar granule cell layer and examined how GCL architecture contributes to granule cell (GrC) combinatorial diversity. We found that GrC combinatorial diversity saturates quickly as mossy fiber input diversity increases, and that this saturation is in part a consequence of short dendrites, which limit access to diverse inputs and favor dense sampling of local inputs. This local sampling also produced GrCs that were combinatorially redundant, even when input diversity was extremely high. In addition, we found that mossy fibers clustering, which is a common anatomical pattern, also led to increased redundancy of GrC input combinations. We related this redundancy to hypothesized roles of temporal expansion of GrC information encoding in service of learned timing, and show that GCL architecture produces GrC populations that support both temporal and combinatorial expansion. Finally, we used novel anatomical measurements from mice of either sex to inform modeling of sparse and filopodia-bearing mossy fibers, finding that these circuit features uniquely contribute to enhancing GrC diversification and redundancy. Our results complement information theoretic studies of granule layer structure and provide insight into the contributions of granule layer anatomical features to afferent mixing.

**Significance Statement:** Cerebellar granule cells are among the simplest neurons, with tiny somata and on average just four dendrites. These characteristics, along with their dense organization, inspired influential theoretical work on the granule cell layer (GCL) as a combinatorial expander, where each granule cell represents a unique combination of inputs. Despite the centrality of these theories to cerebellar physiology, the degree of expansion supported by anatomically realistic patterns of inputs is unknown. Using modeling and anatomy, we show that realistic input patterns constrain combinatorial diversity by producing redundant combinations, which nevertheless could support temporal diversification of like-combinations, suitable for learned timing. Our study suggests a neural substrate for producing high levels of both combinatorial and temporal diversity in the GCL.

## Introduction

Expansion recoding is a leading hypothesis for the role of the dense cerebellar granule cell layer (GCL) (Marr, 1969; Albus, 1971). This network consists of vast numbers of small granule cells (GrCs) that possess on average just four dendrites (Eccles et al., 1967; Herculano-Houzel, 2010). Inputs to this layer are dominated by mossy fiber rosettes (MFRs), which are large presynaptic terminals that branch off of mossy fiber axons (MFs) and convey sensorimotor information from numerous structures into the cerebellum. MFRs form the core of synaptic glomeruli where they are contacted by numerous GrC dendrites, such that each GrC samples around 4 MFRs and sparsely represents convergent afferent input in higher dimensional space, which is thought to be critical for sensorimotor integration in service of motor learning (Fig. 1A; Marr, 1969; Albus, 1971; Blomfield and Marr, 1970). Many studies support these ideas, and similar anatomical organization has been observed in brain areas as diverse as electric fish electrosensory lateral line lobe, fruit fly mushroom bodies, mammalian olfactory cortex and dorsal cochlear nucleus, suggesting a conserved computational function (Kennedy et al., 2014; Sawtell, 2010; Caron et al., 2013).

**Figure 1.**
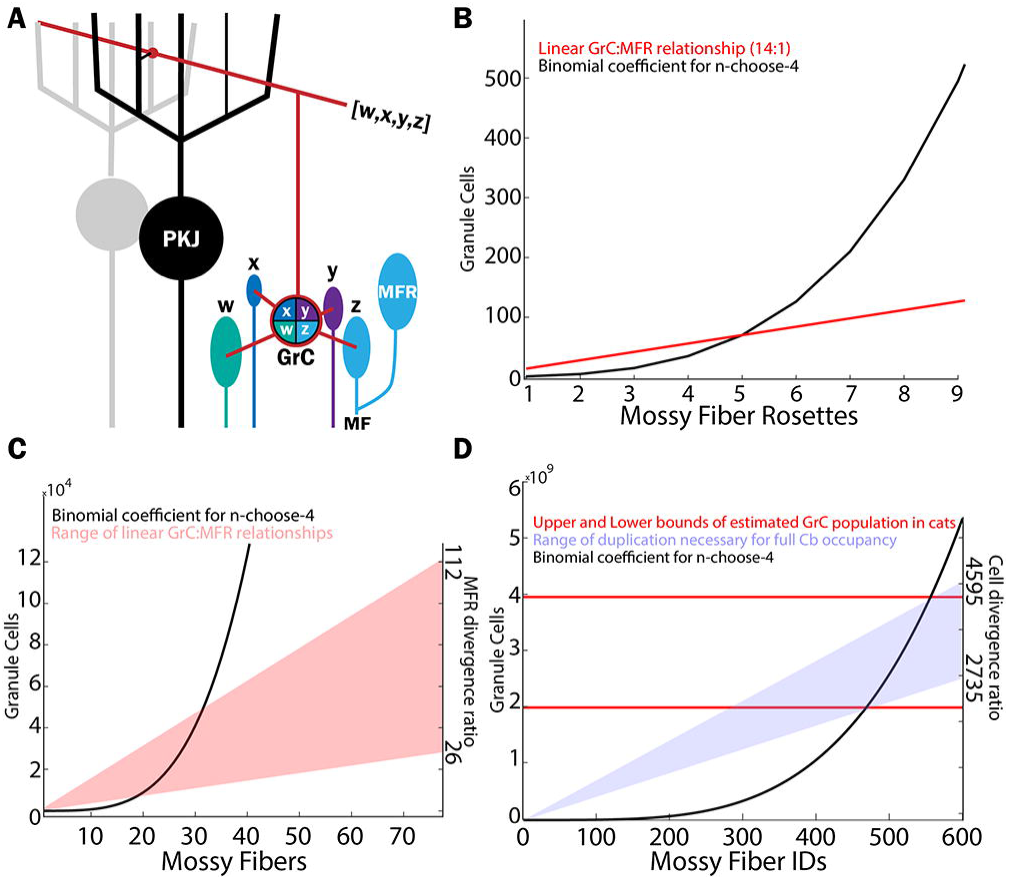
Theoretical limitations of granule cell input combinations based on mossy fiber rosette (MFR)-to-granule cell (GrC) ratios. **A.** Schematic diagram illustrating the MFR-GrC-Purkinje circuit and terminology. MFRs with identities (IDs) w, x, y, z, converge onto a specific GrC. This combination of MFRs to this GrC represents the “quartet” for the illustrated GrC. Throughout the study, we examined how the diversity of MFR combinations onto GrCs relates to patterns of MFR inputs. B. The number of theoretical permutations of MFRs is given by the binomial coefficient, defined by n choose k with replacement, where n is the number of MFRs and k is the number of inputs combined per GrC. The binomial coefficieant, for n choose 4 with replacement (black), is overlaid with the linear function (red) showing the ratio of GrCs to MFRs. The intersections of the curves indicate that based on the linear relationship of MFRs to GrCs, GrCs could fully permute a maximum of 5 unique MFRs. **C.** Similar to (B), but with the linear function relating GrCs to mossy fibers (MFs), taking into account multiple MFRs per MF axon. **D.** The binomial coefficient (black trace; n choose 4 with replacement) is plotted against the number of unique MF IDs that could theoretically be permuted (equivalent to n). The cat cerebellum contains an estimated 2-4 billion GrCs (red lines), indicating that the cat GCL could fully permute 470 to 560 different MFRs. To distribute those MFR identities over the 77 million MFRs estimated to occupy the cat cerebellum, these unique sources would have to be duplicated between 2735-4595 times. Abbreviations: GCL, granule cell layer; MF, mossy fiber; GrC, granule cell; PKJ, Purkinje; MFR, mossy fiber rosette.

A central tenet of the recoding hypothesis is that GrC activity is very sparse, owing to the extensive combinatorial diversity of GrC inputs, where the likelihood of two GrCs sharing the same combination of MFR inputs is low. Yet recent studies using Ca^2+^ imaging to monitor GrC population activity have called into question the sparseness of GrC activity. These studies have noted higher densities of active GrCs than predicted by classic theory (Giovannuci et al., 2017; Wagner et al., 2017; Knogler et al., 2017). How dense activation of GrCs would be supported by highly diverse combinations of inputs onto GrCs is unclear. Furthermore, these dense activity patterns would seem to suggest degradation of the high dimensionality produced by sparse GrC activity, raising the question of the computational utility of redundant GrCs.

While Marr’s original study assumed relatively uniform access of GrCs to mossy fiber afferents, precerebellar sources encoding diverse signals often ramify in dense patches, suggesting non-uniform mixing of inputs. Such anatomical features have contributed to refinement of cerebellar cortical theory (D’Angelo, 2017; Billings et al., 2014). For instance, spatial correlations of MFR inputs can enhance information transmission in models of the GCL (Billings et al., 2014). Furthermore, studies of delay eyelid conditioning reveal that well-timed learning occurs even with dense electrical activation of MFs (Steinmetz et al., 1986; Freeman and Rabinak, 2004, Halverson, 2009). These observations have led to theories proposing temporal expansion of GrC population activity, where GrCs receiving similar inputs nevertheless sparsely fire throughout the conditioning window (Medina et al., 2000; Mauk and Donegan, 1997).

We propose that dense GrC activity patterns could be explained by mossy fiber ramification patterns, and that this density could support temporal expansion processes in the GCL. We used an anatomically realistic model to test the hypothesis that dense ramification patterns of MFRs would produce redundant MFR combinations on GrCs, potentially contributing to denser activations than originally proposed. We also examined how other morphological and organizational features of the GCL – namely MF diversity, GrC dendrite length, and a morphological specialization of MFRs composed of long, thin synaptic extensions that contact GrCs, called filopodia – contribute to and constrain GrC combinatorial diversity. Anatomical details in the model were validated in empirical observations of the nucleocortical and pontine MF systems, described here. Together, our findings illuminate both the capacity of the layer to confer mixed selectivity to GrCs in service of pattern separation (Rigotti et al., 2013; Litwin-Kumar et al., 2017), and the level of redundancy (i.e. the number of identical MFR combinations) likely to emerge within the layer in support of temporal expansion encoding.

## Materials and Methods

The goal of this study was to address how anatomical features of the cerebellar granule cell layer (GCL) influence granule cell (GrC) combinatorial diversity, i.e. the uniqueness of mossy fiber rosette (MFR) combinations made by GrCs. To address these questions, we combined anatomical observations with a model that mimics the geometric organization of the GCL. We varied model parameters to test the role of specific anatomical features to GrC combinatorial diversity. Anatomical and modeling methods are described below.

### Anatomy

#### Subjects

Adult C57/B6 mice (Charles River; n = 9 mice) of either sex were used in accordance with the National Institutes of Health Guidelines and the Institutional Animal Care and Use Committee at the University of Colorado Anschutz Medical Campus. Animals were housed in an environmentally controlled room, kept on a 12:12 light/dark cycle and had ad libitum access to food and water. A total of 9 mice were used in the entire study.

#### Virus Injections

For all surgical procedures mice were anesthetized with intraperitoneal (IP) injections of a ketamine hydrochloride (100 mg/kg) and xylazine (10 mg/kg) cocktail, placed in a stereotaxic apparatus and prepared for surgery with a scalp incision. Craniotomies were made above the cerebellar nuclei (CbN; 1 injection from lambda: 2.0 mm posterior, 1.0 mm lateral, 2.5 ventral; n=9/9 mice); and the basilar pontine nuclei (from bregma) 4.0-4.5 mm posterior, 0.4 mm lateral, and 5.5 mm ventral (n = 4/9 mice). Pressure injections of 0.15-0.25 µL AAV1.hSyn1.mCherry (University of North Carolina Vector Core) and AAV1.hSyn1.eYFP (UNC) were made using a 1 µL Hamilton Neuros syringe attached to the stereotaxic apparatus (Stoelting). Virus use was approved by and in accordance with the University of Colorado Anschutz Institutional Biosafety Committee. All surgeries included postoperative analgesia with IP injections of carprofen (5 mg/kg) once per 24 hr for 48 hr. Mice were housed postoperatively for 3-6 weeks before perfusion to allow for viral expression throughout the entirety of the axonal arbor.

#### Tissue Preparation for Light Microscopy

Mice were deeply anesthetized with an IP injection of sodium pentobarbital (Fatal Plus; Vortech Pharmaceuticals), and perfused transcardially with 0.9% saline followed by 4% paraformaldehyde in 0.1 M phosphate buffer (PB). Brains were removed and postfixed for at least 24 hours then cryoprotected in 30% sucrose. Brains were sliced in 40 µm serial coronal sections using a freezing microtome, stored in PB, and coverslipped in Fluoromount-G (SouthernBiotech) mounting medium.

#### Anatomical Measurements

A total of 1658 MFRs in 9 mice were analyzed, spanning cerebellar lobules including Crus I, Crus II, Paramedian, Simple, and vermal lobules 3, 4, 5, and 6. MFRs were labeled from cerebellar nuclear and/or pontine injections. Rosettes were imaged on a Marianas spinning disc confocal microscope with a 63x objective. To quantify nearest neighbors, montages of Lobules 6 and Crus 1 were produced from high resolution images, and the location of each MFR was mapped (n = 874 boutons; n = 4 dual injected mice, Lobules 6 and Crus 1). We noted the presence or absence of filopodia on these MFRs and an additional 784 MFRs from 5 additional mice located throughout the cerebellum for a total of 1658 MFRs. Euclidean distance from each rosette to its 4 nearest neighbors was then computed and analyzed in MATLAB (RRID:SCR_001622). To quantify the number and distance of the filopodia from the MFR, 84 rosettes were analyzed, with 70 fully reconstructed in 3D using Neurolucida 360 software (RRID:SCR_001775) (Fig. 6). Filopodial boutons were defined as swellings at least 1 µm wide on processes that extended from the main rosette but did not leave the section (Gao et al., 2016).

#### Experimental Design and Statistical Analyses

Anatomical observations were made in 4-9 mice. Nearest neighbor measurements were made with custom scripts in MATLAB and compute the Euclidean distances of the 4 nearest neighbors of each MFR in 2 dimensions in the coronal plane. Summary data of nearest neighbors are visualized in cumulative distribution functions and populations compared with the two sample Kolmogorov-Smirnov test in Matlab, with p-values and n’s reported in the text. Simulations were validated by performing multiple instantiations of modeled systems described in each section below.

### Modeling

#### Model-free calculations

Theoretical numbers of MFR combinations given the number of synapses per GrC, were computed using n choose k with replacement, where n is the number of MFRs and k is the number of inputs per GrC (Fig. 1).

#### Simulations

We modeled groups of mossy fibers and granule cells in MATLAB with density and spatial relationships based on anatomical and physiological data (Palkovits, et al., 1971; Sultan et al, 2001; Solinas et al., 2010). Specifically, the model incorporated physiological density and distribution of GrCs and MFRs, the length of GrC dendrites, and the divergence and convergence of MFRs onto GrCs (Table 1).

**Table 1:**
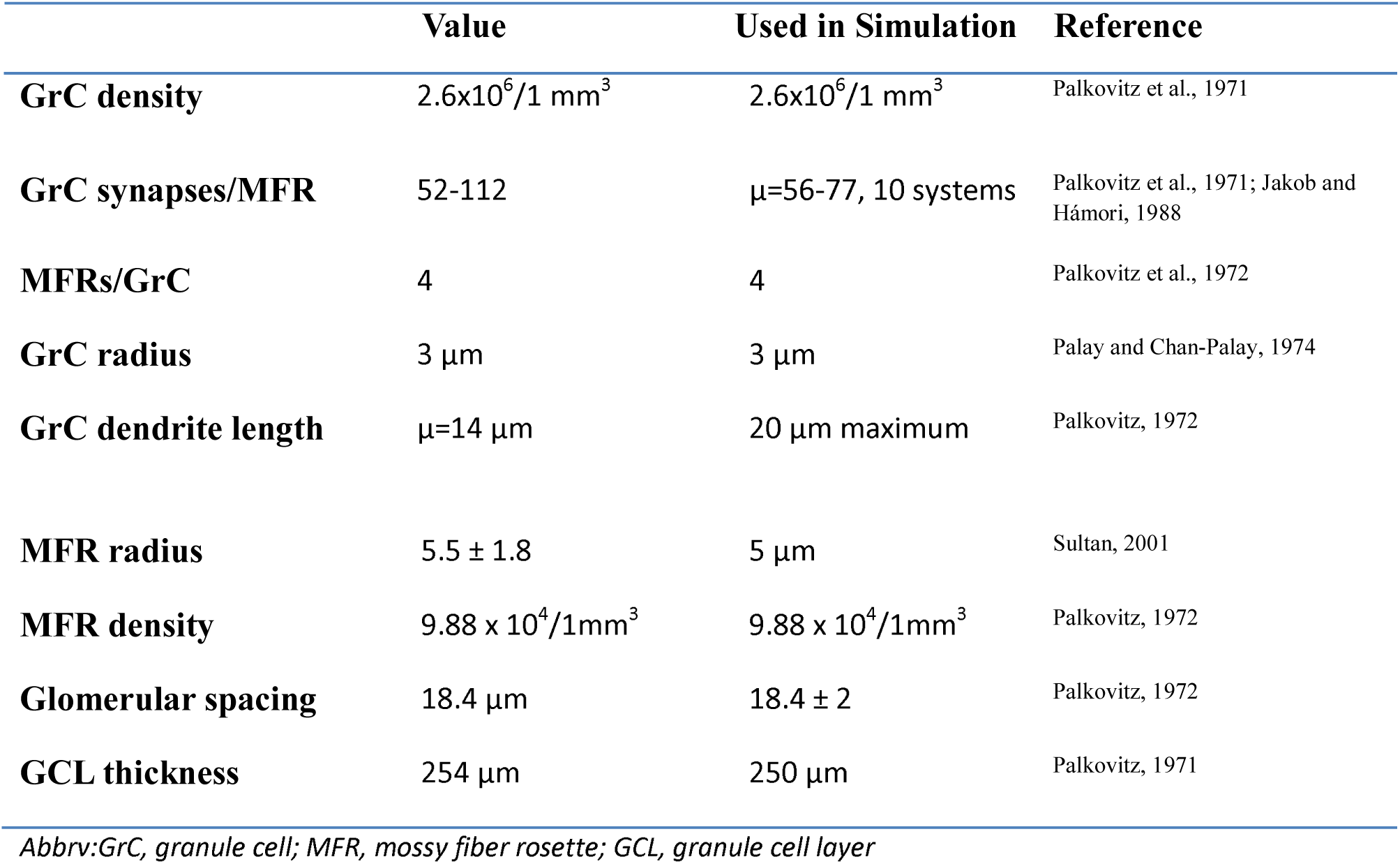
Anatomical Data Used in Simulations. Values used in simulation and 1067 references cited for each.

| Value | Used in Simulation | Reference |
| --- | --- | --- |

|  |  |  |  |
| --- | --- | --- | --- |
| **GrC density** | $2.6 \times 10^6 / 1 \text{ mm}^3$ | $2.6 \times 10^6 / 1 \text{ mm}^3$ | Palkovitz et al., 1971 |
| **GrC synapses/MFR** | 52-112 | $\mu=56-77$, 10 systems | Palkovitz et al., 1971; Jakob and Hámori, 1988 |
| **MFRs/GrC** | 4 | 4 | Palkovitz et al., 1972 |
| **GrC radius** | $3 \mu\text{m}$ | $3 \mu\text{m}$ | Palay and Chan-Palay, 1974 |
| **GrC dendrite length** | $\mu=14 \mu\text{m}$ | $20 \mu\text{m}$ maximum | Palkovitz, 1972 |
| **MFR radius** | $5.5 \pm 1.8$ | $5 \mu\text{m}$ | Sultan, 2001 |
| **MFR density** | $9.88 \times 10^4 / 1 \text{ mm}^3$ | $9.88 \times 10^4 / 1 \text{ mm}^3$ | Palkovitz, 1972 |
| **Glomerular spacing** | $18.4 \mu\text{m}$ | $18.4 \pm 2$ | Palkovitz, 1972 |
| **GCL thickness** | $254 \mu\text{m}$ | $250 \mu\text{m}$ | Palkovitz, 1971 |

*Abbrev: GrC, granule cell; MFR, mossy fiber rosette; GCL, granule cell layer*

MFRs and GrCs were programmed to represent their 3D position in space and associated radii. Model systems were generated by first populating a defined space with MFRs. System dimensions were 100×100×250 µm, and contained 3458 GrCs and 247 MFRs, except where noted. Each MFR was checked against every other existing MFR to ensure spatial non-overlap, rejecting a placement if the distance between the centers was less than the sum of the radii. MFR tiling obeyed a flexible spacing constraint such that each MFR had three neighbors with a mean distance of 18.4 microns between them from center to center, consistent with anatomical measurements of glomerular spacing (Palkovits, et al., 1971). To do this, we chose randomly from existing MFRs and placed a new rosette at 16.4-20.4 µm intervals. Perfectly even spacing of MFRs generated tetrahedrons, which do not tile in 3D space, so up to 2 microns of jitter about the mean spacing allowed for generation of nearly evenly spaced MFRs. MFR tiling approximated a tetrahedral matrix.

GrC placement followed a series of steps. First, seed locations were selected from either existing MFRs or GrCs, then a new GrC was placed a random distance away, excluding locations overlapping within the sum of the radii of existing elements. Second, GrCs made four random synaptic connections with nearby MFRs, limited to distances less than or equal to the sum of the dendrite length and MFR radii (28 µm, unless where noted). If the random location was not within reach of 4 MFRs it was discarded.

Third, GrCs synapsed preferentially onto MFRs that had fewer than 56 synapses with other GrCs, but MFR connectivity was capped at 80 GrCs. Overall, there was a range of 53-77 GrC synaptic connections per MFR, with a mean of 56 (SEM = 0.82, n=151 systems). The final result was a modeled population with closely packed elements resembling the dense packing of GrCs within the GCL (Fig. 2A).

**Figure 2.**
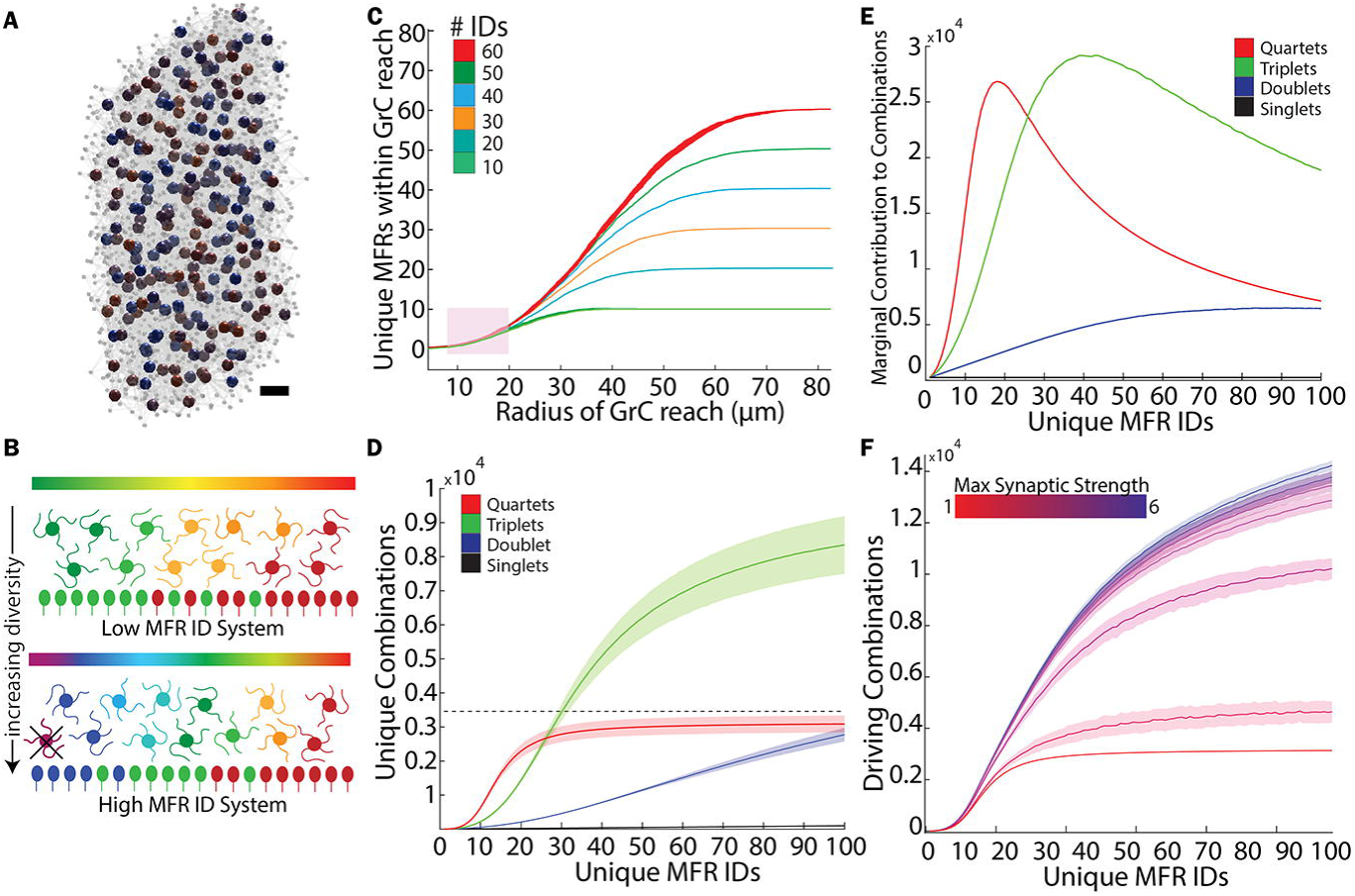
The spatial organization of the granule cell layer (GCL) restricts combinatorial expansion. A. We constructed spatially realistic models of the granule cell layer (GCL), shown here in a simplified rendering. Large spheres represent simulated MFRs, with colors representing ID numbers. Grey spheres represent GrCs, and GrC dendrites are grey lines. A 247 MFR system is depicted here, with all synaptically connected GrCs per MFR rendered. Scale bar is approx. 25 µm. **B.** Cartoons illustrating low and high diversity systems. Bottom rows illustrate MFRs with ID indicated by color. GrCs (middle), combine local MFR inputs to assume a unique combinatorial identity (blended colors). Throughout the remainder of the study, we examine how MFR diversification and clustering in a spatial model of the GCL influences GrC combinatorial diversity, testing, for example, how spatial segregation of inputs favors particular combinations (e.g. the purple GrC, indicating a theoretical combination of red and blue MFRs cannot exist, because red and blue MFRs are too far apart). **C**. Effect of GrC dendrite length on access to unique mossy fiber rosettes (mean +/- SEM; n = 10 systems with 10 trials each, for each MFR diversity level). An anatomically realistic dendritic range between 12-20 μm restricts access to roughly 6 unique inputs, regardless of the diversity of the system. **D**. Populations of combinatorially unique GrCs as defined by their k-input combinations (quartets, triplets, etc.) as a function of increasing numbers of unique MFR identities uniformly distributed across the population. Dashed line indicates the total number of GrCs in the system. **E**. A function plotting the relative contribution of each additional MFR ID to the total number of MFR combinations made by GrCs (Eq. 1). As the model diversifies with increasing numbers of unique MFRs, fewer new combinations are produced per additional MFR ID number, peaking at 5-20% diversity (i.e. between 5 and 20 MFRs/100 share the same ID) for quartet and triplet systems. This peak indicates the point beyond which increasing diversity contributes relatively less to the number of unique MFR combinations made by GrCs. **F**. The effect of MF long term potentiation on combinatorial representation. With a GrC firing threshold of 4 active inputs, a full quartet of active MFRs is required to drive activity. As the strengths of inputs increase, the mean number of potential combinations that can drive a given GrC increase.

#### MFR diversity and spatial distribution modeling

One goal of the present study was to understand how MFR diversity influences GrC combinatorial diversity. Because we were interested in identifying specific combinations of MFRs onto GrCs, we assigned an identity “ID” to each MFR. The ID terminology is a proxy for the source and/or uniqueness of the MFR, such that we can think of modeled MFRs as originating from different cells or different nuclei if they do not share an ID number. Throughout the text we refer to MFRs as a general term that assumes each MFR has an ID, such that groups of MFRs imply the specific combination of ID numbers of MFRs converging on a GrC.

To examine the relationship between MFR diversity and GrC combinatorial diversity, we varied the number of different ID numbers assigned to a fixed number of MFRs. We refer to the number of different ID numbers in a system as *n*. In low diversity systems, many MFRs had the same ID number, whereas in high diversity systems, few to no MFRs shared an ID. We analyzed the combinations of MFR ID numbers formed by GrCs within the simulation, tracking the number of shared ID number combinations between different GrCs. The combinatorial diversity of GrCs refers to the number of different MFR combinations produced by the population of GrCs within the simulation. In some analyses, groups of MFRs converging on a GrC were subdivided into the quartet, (the combination of 4 MFRs), triplets (any 3 of the 4 convergent MFRs), etc., noting that these MFR combination sizes are equivalent to the “codon” terminology used in Marr, 1969. We also investigated the relative change in GrC combinatorial diversity produced by adding one additional ID number to the MFRs in the system, defined as,

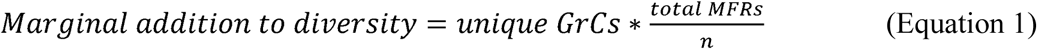

where unique GrCs refers to the number of unique MFR ID number combinations produced by the system GrCs, total MFRs is the number of MFRs in the system, and *n* is the number of ID numbers assigned to the MFRs.

Another feature of mossy fiber patterning that we simulated was clustering of similar MFRs. To examine the effect of MFR clustering on the diversity of MFR combinations produced by GrCs, we populated the simulation with MFRs assigned ID numbers drawn from a Gaussian-like probability distribution generated using the randn function in MATLAB. In effect, this method led to non-uniform representation of specific ID numbers within the system, with many MFRs assigned highly probable ID numbers, mimicking clustering. These systems were compared to non-clustered models, in which ID numbers were selected from a flat ID probability distribution, producing a space in which each ID number was equally likely. These systems were analyzed for diversity, redundancy and fraction of theoretically possible combinations produced, with analyses repeated in 100 trials across 100 systems (120 × 120 × 100 µm in size, containing 142 MFRs and 1,988 GrCs). Redundancy of a particular combination of MFRs on modeled GrCs was defined as the number of GrCs in that system possessing identical combinations of MFRs (i.e, the same quartet). The system redundancy was the mean number of repeats of MFR combinations formed by GrCs in that system. Diversity was defined as the total number of unique combinations of MFR inputs of a given size in the GrC population. We refer to redundancy changes as a function of MFR diversity as redundancy(*n*) where *n* is the number of different ID numbers.

A related spatial pattern of MFR termination that we modeled was sparsity. Here, a random MFR was chosen from a standard model and was given a new, unique ID number. This analysis was repeated across a range of ID diversity levels in 150 different system instantiations with 5 trials each. To analyze the diversity and redundancy of MFR combinations with sparse fibers included, three different types of systems were compared. First, systems with ‘*n*’ inputs, called “baseline” systems, where ‘*n*’ is the number of different ID numbers; Second, systems with ‘*n*’ inputs, in which a single rosette is changed to a unique input, called “sparse” systems; Third, systems with ‘*n*+1’ inputs distributed uniformly, termed “expanded” systems. The expanded system was essential to the comparisons because it allowed us to determine if the effect of sparseness on GrC combinatorial diversity was solely a consequence of adding an additional input. Using these systems, we first calculated differences in diversity and redundancy in the sparse and expanded systems relative to baseline. We then compared changes in diversity and redundancy between these conditions, expressed as a percent difference in the effect of the sparse or uniformly expanded inputs, defined by:

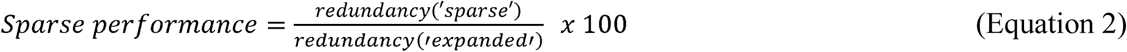

The diversity measurements were identical, substituting redundancy for diversity.

We postulated that dense afferents serve a computational purpose that could be disrupted by adding another uniformly distributed input. We therefore asked whether a sparse input, when added to a pre-existing system, preserves GrC combinatorial patterns differently than adding another uniform input. We analyzed the retention of existing GrC combinations in the baseline system upon addition of a “sparse” or “expanded” input type by computing a retention index: First, each MFR quartet in the baseline system was given a value of 1, with redundant MFR combinations summed, such that the value reflected the number of repeats of a given combination. Next, each combination in the ‘sparse’ or ‘expanded’ systems was scored similarly. Finally, each combination in the baseline system was then compared to each combination in the new system, receiving a retention score given by baseline value divided by new system value. If a combination was lost in the new system, it was scored as a 0. The mean score for all combinations was computed for comparisons between baseline and ‘sparse’ and baseline and ‘expanded’. Finally, to compare how different types of systems retained combinations we computed a metric that took the ratio of retention scores between the systems, defined as:

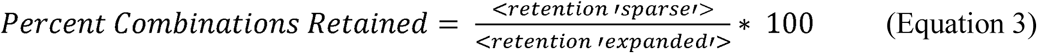

#### Dimensionality Calculations

The GCL is hypothesized to act as a combinatorial expander, increasing the dimensionality of inputs (Marr, 1969; Albus, 1971). Because dimensionality is a function of the independence of neuronal activity, coincident neural activity produced by redundant MFR combinations would be expected to reduce dimensionality. Therefore, to explore the dimensionality of systems as a function of MFR diversity, we generated 150 different systems of GrCs and MFRs as described above. Within each system, we simulated a range of MFR diversity levels where between 1-100 different MFR ID numbers were assigned to the MFR population. We established a proxy for neuronal activity where GrCs were considered active when three of their four MFRs were synchronously active (Billings et al., 2014; Jörntell and Ekerot, 2006). The MFR activity level was set such that on average 10% of MFRs in the simulation were active at a given time (Litwin-Kumar et al., 2017), over 1000 epochs. The activity of a given MFR ID number was determined by assigning it a value from a random number generator every epoch. When the random number exceeded a pre-determined threshold, the ID number was ‘active’. Note than in low diversity systems, many MFRs share the same ID number, rendering many MFRs coincidently active. The threshold for determining an active ID number from the random number generator was therefore varied to ensure that even in low diversity systems, with many MFRs possessing the same ID, the mean total active population was maintained at 10%. 1000 epochs were generated in each simulation and all pairwise GrC correlations and variance of correlations were measured over the all epochs. 5 trials of each simulation were performed and the average correlation and variance measured and used in Equation 4 (below). In total, 500,000 epochs for each diversity level were analyzed.

To assay the dimensionality produced by combinatorial expansion (CE) of the model GCL as a function of diversity level, *n*, we used the equation,

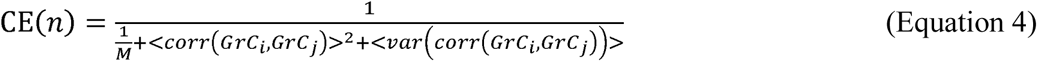

where n is number of unique MFR ID numbers in the model, M is the number of granule cells, <corr(GrCi,GrCj)>^2^ is the squared mean correlation coefficient of the activities of GrCi and GrCj over all pairwise GrC comparisons, *i* and *j*, across all epochs, and <var(GrCi,GrCj)> is the mean variance of all pairwise correlations across all epochs as described in Litwin-Kumar et al., 2017. This calculation was repeated for each system and averaged for each value of *n*.

A prominent hypothesis in cerebellar literature proposes that GrCs receiving correlated inputs become temporally diversified (i.e. fire at different times despite receiving the same input), leading to increased dimensionality supporting learned timing (Fig. 3A,C; Mauk and Donegan, 1997; Medina et al., 2000), which is not captured in Equation 4. Behavioral measurements of intervals over which an animal can learn a conditioned response suggest that information can be diversified up to 500 ms, with physiological measurements showing individual GrCs bursting for approximately 20 ms (Schneiderman and Gormezano, 1964; Smith et al., 1969; Yeo and Hesslow, 1998; Ishikawa et al., 2015). At the limit, every GrC could be decorrelated from every other GrC, producing maximal dimensionality even if each GrC received identical inputs. However, based on GrC burst times, this would predict unrealistically long temporal expansion windows. We therefore used physiological estimates of temporal learning windows to constrain a metric for temporal expandability (TE) where we penalized over- or under-representation of dimensions produced by CE, for a given time window. We defined ‘over-representation’ (R_o_) and under-representation (R_u_) as:

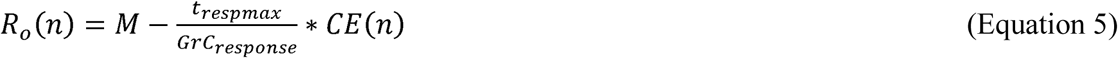

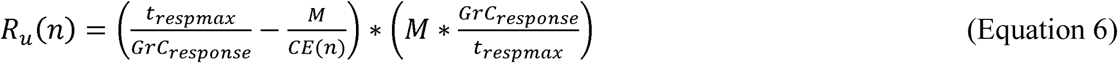

**Figure 3.**
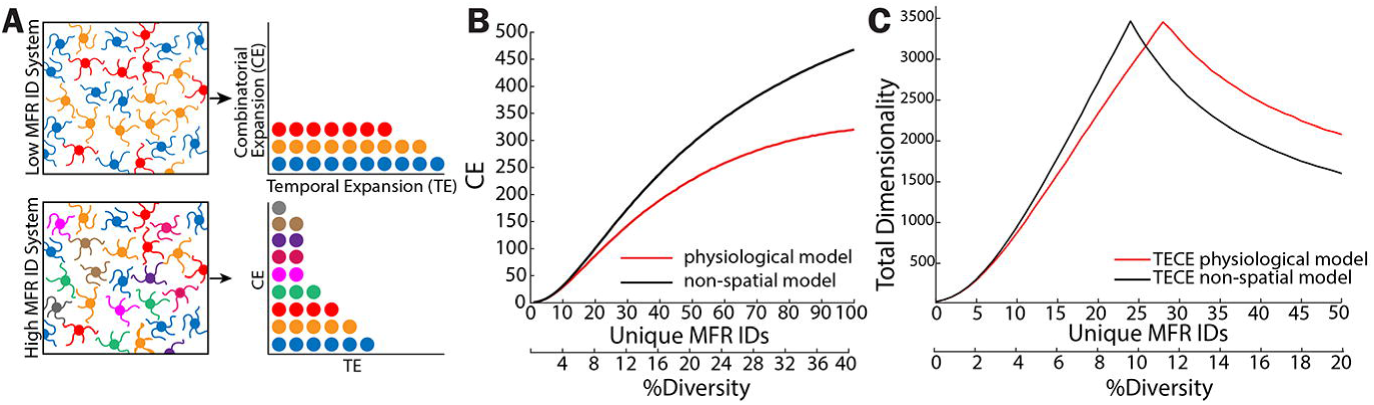
Spatial patterning enhances temporal expansion of GrC representation. **A**. Cartoon illustrating the hypothesized tradeoff between combinatorial expansion (CE) and temporal expandability (TE), which requires redundant MFR combinations (shared colors) by multiple GrCs. **B**. Combinatorial expansion dimensionality plotted as a function of MFR diversity for the spatially-constrained model “physiological” and non-spatial model. **C**. The composite of combinatorial expansion and temporal expansion (TECE) for physiologically constrained (red) and non-spatial (black models) for a time window of 500 ms. At higher MFR diversity levels, the spatially realistic model possesses greater TECE dimensionality than the non-spatial model because it both expands combinations and produces redundancy required for temporal expansion of dimensions.

where *n* is the number of different MFR ID numbers, M is the number of GrCs and CE(*n*) is the dimensionality of the system computed using Eq. 4. The R_o_ formula returns the number of “extra” GrCs over all dimensions that are unnecessary for complete temporal expansion over the t_respmax_ interval (500 ms), assuming a given GrC_response_ burst duration (20 ms). The R_u_ formula returns the difference between the number of GrCs that would be needed to completely represent the time window and the number of GrCs per dimension, scaled by the optimal number of GrCs needed per dimension to represent the time interval. When either R_o_ or R_u_ returned a negative value, we set the metric to 0.

We then used these penalties to compute a function that captures both the temporal and combinatorial expansion (TECE):

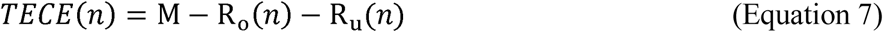

Where M is the number of GrCs (and maximum dimensionality), from which the lost dimensionality which occurs due to the R_o_ and R_u_ terms are subtracted.

Finally, we tested differences in the TECE(*n*) calculated for models in which spatial relationships were included (‘physiological’) and models in which there were no spatial constraints considered (‘non-spatial’). The non-spatial model used here was simulated by assigning 4 MFR IDs at random to each GrC in a model system of the same size as the ‘physiological system’, irrespective of space.

#### Filopodia Modeling

Statistics of MFR filopodia gathered from the anatomical experiments were used in the spatial GCL model to mimic fiolopodia. We simulated filopodia by adding between 1-5 (median, 2) synaptic connections with GrCs within 22 microns of the MFR. We added simulated filopodia to between 8 - 20% of MFRs. This was simulated in 450 different systems, with 5 trials each.

## Results

### Theoretical limits on combinatorics imposed by granule cell-to-mossy fiber ratios

Expansion recoding is a leading hypothesis for the role of the dense cerebellar granule layer network formed with mossy fiber rosettes (MFRs). First proposed by Marr and elaborated by Albus (Marr, 1969; Albus, 1971), the idea that granule cells (GrCs) sample around 4 MFRs and represent convergent afferent input in higher dimensional space is central to ideas of cerebellar sensorimotor integration (Fig. 1A). Before analyzing how GCL organizational features influence the number of different combinations of MFRs represented by the GrCs, i.e. combinatorial diversity, we made a series of simple calculations defining the maximum number of MFR permutations possible, given MFR diversity. These calculations highlight the fact that complete permutation of cerebellar inputs is not physiologically realistic and motivate the spatially-constrained modeling. The theoretical number of permutations of MFRs can be computed using the binomial coefficient, n choose k, with replacement, where n is the number of unique MFRs and k is the number of inputs per GrC. We compared these values to the number of GrCs that exist per MFR (14:1), based on numerical ratios derived from anatomical estimates (Jakob and Hamori, 1988). The rapid increase in the binomial coefficient as a function of MFR numbers quickly exceeds the number of GrCs that exist (Fig. 1B-C) such that when there are more than 5-6 MFRs in a system of 84 GrCs, some MFR quartets go unrepresented in the GrC population (Fig. 1B). Even when considering divergence, where each MF forms between 20-200 MFRs (Wu et al., 1999; Shinoda et al., 1992; McCrea et al., 1977; Quy et al., 2011), theoretical complete mixing is limited to between 18 and 31 unique sources in the GrC population of approximately 6,000-46,000, depending on the level of divergence (Fig. 1C, intersection of shaded regions and black curve). These calculations indicate that the cat granule layer, containing 2-4 billion granule cells and 77 million MFRs (Palkovitz et al., 1971), could fully permute just 470-560 unique MFR sources, depending on the level of MF divergence. Therefore, the cerebellum would not represent all quartets unless each input is duplicated between 2,700-4,600 times, a highly unlikely scenario (Fig. 1D, blue shading).

### Spatial constraints on granule layer combinatorics

The previous calculations do not take into account any spatial features of GCL organization, which would be expected to influence the diversity of convergence of MFRs onto GrCs. As an extreme example, two MFs that terminate in different lobules would obviously not converge on a common GrC. We speculated that the morphological features of the GCL might extend these spatial restrictions at local levels, where combinations of MFs ramifying even in a relatively local area might not be combined if GrC dendrites are too short to reach them (schematized in Fig. 2B). We therefore examined how the anatomy of the GCL influences the performance of the layer as a combinatorial expander. To address this, we developed a model that mimics the geometric organization of the GCL and manipulated spatial features to determine the role of specific anatomical features to GrC combinatorial diversity (Fig. 2A; See Modeling in Materials and Methods).

We first investigated the role of GrC dendrite length on access to local MFRs. We measured the effect of GrC dendrite length on the access to diverse MFRs in modeled systems with varying MFR diversity. We found that dendrite lengths falling within a physiological range (8-20 µm; shaded region) strongly limit access to MFRs. As the MFR diversity increases, GrCs cannot access that diversity because of the shortness of their dendrites (Fig. 2C). Therefore, locally, the number of MFRs accessible to granule cells is capped at fewer than 10, regardless of MFR heterogeneity. As expected, lengthening dendrites permitted granule cells to access a greater diversity of MFRs. Increasing the dendrite length from 20 to 60 microns allows a GrC to access 5-10-fold more unique MFRs, depending on the diversity of the local MF population (Fig. 2C).

We next computed the number of unique combinations of 4 MFRs produced by a modeled population of ~3500 granule cells and ~240 MFRs, taking into account spatial relationships of GrCs and MFRs. We assigned ID numbers to MFRs in the model and varied the diversity of the population of MFRs from being entirely homogeneous (where each MFR has the same ID) to entirely heterogeneous (where each MFR has a unique ID; Fig. 2D). Focusing here first on quartets (combinations of 4 inputs), the relationship between MFR diversity and number of different combinations formed on GrCs showed two striking features. First, GrCs are quickly saturated with unique MFR combinations as MFR diversity increases, illustrated by the asymptote in the number of unique granule cell combinations (Fig. 2D, red curve). This cap on unique MFR combinations suggests that with just moderate MFR diversity, most GrC input combinations are unique. As the MFR population continues to diversify, new unique MFR combinations replace other unique MFR combinations, illustrating the effectiveness of the GCL as a combinatorial expander. The apparent cap on diversification of quartets at approximately 30 MFR IDs in this system indicates that quartets quickly approach the combinatorial limit imposed by GrC population sizes.

A second feature of GrC combinatorial diversity as a function of MFR diversity was that GrC combinatorial diversity is never maximal, asymptoting near 80% of GrCs bearing unique MFR combinations. That is, the number of unique combinations remains below the number of GrCs in the system, even when the MFR population is completely diverse. This phenomenon reflects local resampling of MFRs, and indicates that multiple representations of specific combinations, while not dominant in the population, is a byproduct of short dendrites and glomerular presynaptic structures.

We next examined how the number of GrCs with different MFR combinations changes if we considered just a subset of the combination of 4 convergent MFRs (3, 2, or 1 MFR), since a subset of MFR afferents could produce redundant GrC activity (Billings et al., 2014; Jörntell and Ekerot, 2006). We found that as the diversity of MFRs increases, the number of combinations of 3 MFRs produced by GrCs continues to increase beyond the point where quartets are saturated (Fig. 2D, green curve).

Regardless of combination size, we noticed a roll-off in the number of unique MFR combinations produced as the system diversifies. We explicitly computed the relationship between GrC combinatorial diversity and the addition of each new MFR, plotted in Fig. 2E, by weighing each additional GrC-MFR combination against the ratio of total MFRs to unique MFRs (Eq. 1). The peaks in these curves indicate the point at which each additional MFR ID adds relatively less to overall GrC combinatorial diversity.

These analyses assume equal strength of MFR inputs to GrCs and spiking thresholds requiring coincident activity of either quartets, triplets or doublets. Interestingly, long-term potentiation (LTP) at the MFR->GrC synapse has been observed, which might be predicted to change the effective subset of convergent MFRs onto a granule cell. To test this idea, we established a thresholding rule on a granule cell, but then increased the contribution of each MFR, simulating LTP at this synapse (Mapelli and D’Angelo, 2007). When inputs were allowed to strengthen, we found an increase in the number of combinations that could drive this population of GrCs (Fig. 2F). This diversity in driving inputs might be exploited by an adaptive filter via the Golgi cell (Billings et al., 2014). Furthermore, it would be predicted to produce denser recruitment of granule cells than in an unpotentiated state (Diwakar et al., 2011).

### Spatial constraints enhance the capacity for temporal diversification of granule cells

These data show that the spatial organization of the granule layer limits the number of MFR permutations possible and produces redundant MFR combinations. The presence of redundant combinations of MFRs on GrCs in the spatially constrained model raised the question of whether there may be utility to redundancy that is not well captured by dimensionality produced by combinatorial expansion (CE). Theories of the expansion of specific information in the temporal domain (Mauk and Donegan, 1997; Medina et al., 2000) could explain the utility of redundant combinations, since identical information sources could activate GrCs, which could be further diversified in time, in support of learned timing. For instance, if many GrCs share a dimension produced by combinatorial identity (Fig. 3A, “Low ID system” indicated by color), that dimension could be temporally expanded (TE). Alternatively, if GrCs are already extremely combinatorially diverse, and few GrCs are shared per dimension, the combinatorially defined dimension cannot be temporally expanded as extensively (Fig. 3A, bottom).

These features would predict reduced dimensionality of information represented by the system. We used a simplified model of GrC activity based on MFR activity and analyzed dimensionally as a function of MFR diversity (See Materials and Methods, Eq. 4). We calculated the dimensionality of the system with and without spatial constraints and found that, consistent with recent findings, the anatomically constrained “physiological” system showed a decrease in dimensionality relative to a non-spatial, randomly connected network (Fig. 3B; Litwin-Kumar et al., 2017). We therefore considered the idea that GrCs with redundant combinations of MFRs could fire at different times, owing to either GCL circuitry or synaptic diversity (Medina et al., 2000; Chabrol et al., 2015). This scenario would both recover dimensionality lost by redundancy and support specific information being represented over extended time windows. Based on the temporal dynamics of GrCs and the limit of learned timing in delay eyelid conditioning, we modeled GrC activity such that each GrC was active for a 20 ms epoch (GrC_response_) over a 500 ms window (t_respmax_) (Ishikawa, et al. 2015, Schneiderman and Gormezano, 1964; Smith et al., 1969; Yeo and Hesslow, 1998). We assumed a cost to over- and under-representing a given combination over time (Eq. 5–6, Methods) and measured the dimensionality of the time expanded simulation (TECE) over a range of MFR diversity levels (Eq. 7). Fig. 3C shows the results of this simulation and analysis when added to the calculation of dimensionality produced by combinatorial expansion. Intuitively, the more redundant the combinations, the greater the capacity of the specific combinations to diversify over time. This intuition is supported by a peak in the temporal expansion (TECE) curve at low diversity levels, which then dropped off with increasing MFR diversity. As the diversity of inputs increases, the layer falls short of providing enough GrCs to expand this information in time.

Importantly, the model with physiological spatial constraints showed improved performance over the non-spatial connectivity model, achieving both high dimensionality and temporal expandability (Fig. 3C). The peaks of these curves occur when the number of GrCs per dimension is equal to the minimal number of GrCs required to completely fill the time interval (See methods) and is therefore dependent on the temporal assumptions (GrC_response_ and t_respmax_; Eq. 5–6). The physiological model has higher TECE than the non-spatial model with temporal expansion windows greater than 200 ms, within the range which rabbits readily learn conditioned stimuli (Schneiderman and Gormezano, 1964). At shorter t_respmax_ (i.e. < 200 ms), the non-spatial model has higher TECE. These calculations suggest that the granule layer may balance requirements to expand information temporally as well as diversify inputs as a result of combinatorial expansion.

### Impact of mossy fiber heterogeneity on spatial organization of GrCs sharing inputs

Receptive fields of GrCs tend to be somatotopically patchy, suggesting redundant mossy fiber input to GrCs (Welker et al 1984; Jörntell and Ekerot, 2006; Voogd and Glickstein, 1998). Because MFRs are sampled by many neighboring GrCs, local sharing of a given MFR is expected, and we therefore examined the spacing of GrCs that share a given number of MFRs in our model. We measured the spatial relationship between GrCs that shared one or more common inputs (i.e. inputs had common MFR ID numbers). As MFR diversity increased, the distance between GrCs sharing inputs diminished rapidly (Figs. 4A, B). We analyzed this phenomenon by measuring the Euclidian distance between all GrCs and classified them into groups determined by the similarity of their combinations. For example, with only five MFR ID numbers assigned to the MFR population, the spatial distribution of GrCs that share four specific MFR inputs or share no inputs are equally spaced, illustrated in a cumulative distribution of the distances between GrCs sharing inputs (Fig. 4A). By contrast, when the MFR population is diversified to the point that it is approximately 25% diverse (1 in 4 MFRs shares an ID), GrCs that share the same 4 input ID numbers cluster within approximately 20 microns of one another and GrCs that share no inputs remain homogeneously spaced within the volume (Fig. 4B, C). This can be seen comparing the red and black curves in Fig. 4B, which plots distances of GrCs that share all or no inputs, respectively. These analyses were extended for modeled systems in which we varied MFR diversity systematically, ranging from all identical to all dissimilar. Distances between GrCs with shared input combinations drop with increased MFR diversity (Fig. 4C).

**Figure 4.**
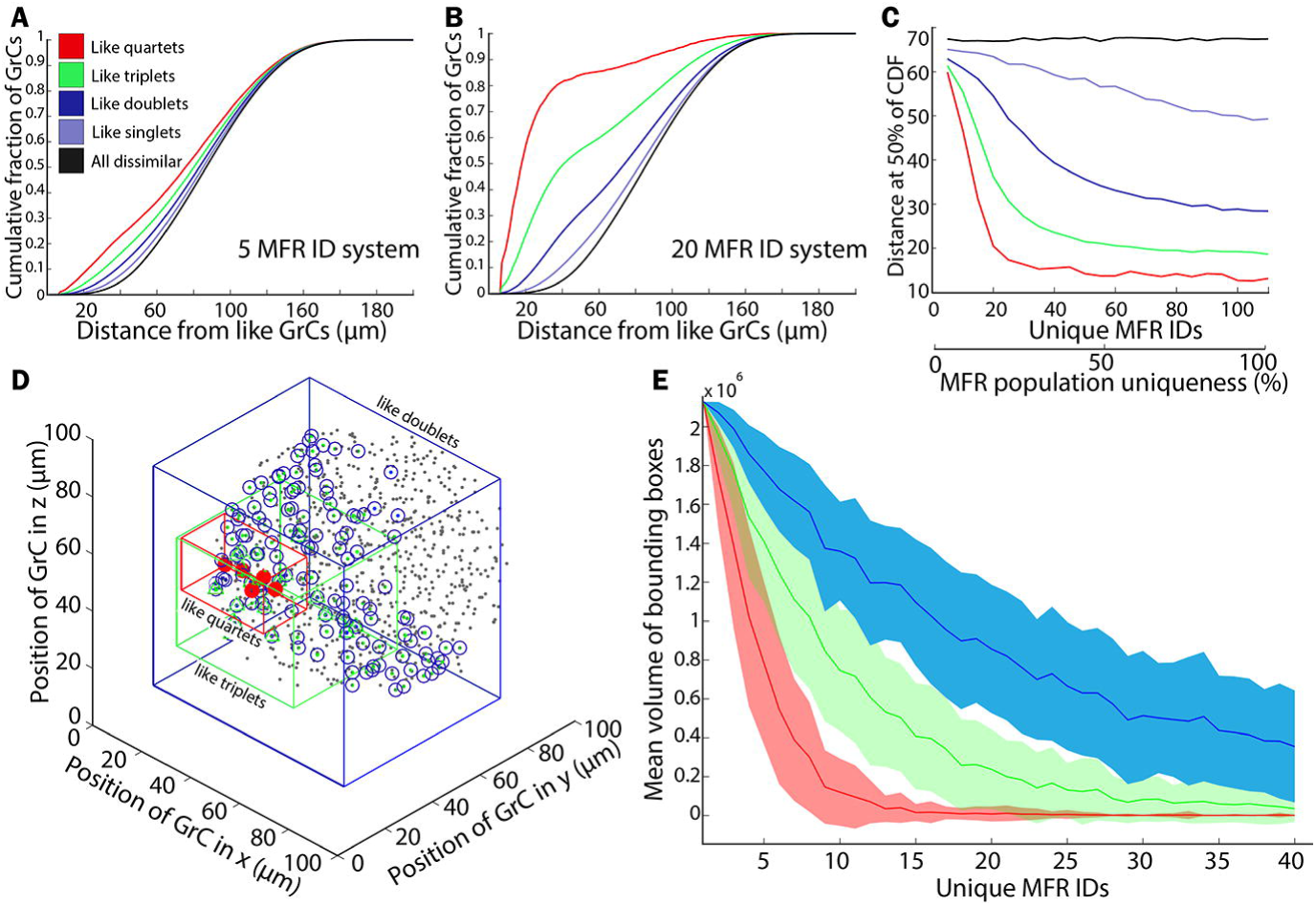
Spatial clustering of GrCs with similar combinations emerges as a consequence of MFR access and diversity. **A.** Cumulative distributions plotting the distance between a GrC and every other GrC in the model that shares a given number of its MFR inputs (i.e. GrCs that share quartets, GrCs that share triplets etc.) in a system with 5 different mossy fibers (5 ID) **B.** Same as (A) but in a system with 20 different mossy fibers (20 ID). Colors as in A. **C.** Summary of clustering effect as a function of MFR diversity, illustrating that dense spatial restriction of like-combinations is most pronounced with high MFR diversity. Colors as in A. **D.** Illustration of spatial extent of GrCs sharing 4 inputs “quartets” (red), triplets (green) and doublets (blue). **E.** Summary of volumes occupied by like-quartets, triplets and doublets as a function of the diversity of MF IDs in the system.

While redundant quartets become highly restricted in space, GrCs that share 3, 2 or 1 MFR ID can remain fairly distant from one another, as shown in Figs. 4B,C (green and blue curves). We visualized the space of shared MFR IDs in a modeled system with 20 MFR ID numbers (Fig. 4D). GrCs are plotted as gray points, and colored boxes surround GrCs that share MFR ID numbers. The volume occupied by GrCs with identical quartets (red box) is extremely circumscribed. The volume of the bounding box for identical quartets drops rapidly as MFR diversity increases (Fig. 4E, red curve), asymptoting around 36 µm^3^, the volume occupied by the reach of a single mossy fiber. Similarly, bounding volumes of GrCs sharing MFR triplets or doublets also decrease as diversity increases (Fig. 4E, green and blue curves), implying that although specific GrC MFR quartet combinations saturate quickly, the number of shared MFRs in the population continues to drop as MFR diversity increases, reducing the space over which GrCs share MFRs.

In summary, by increasing diversity of mossy fibers, patchy representations emerge because of the spatial restrictions of the GrC dendrites and MFRs. Grouping also occurs as a consequence of MFR combination probabilities dropping off sharply in space, although subsets of inputs are shared by more widely spaced GrCs. This phenomenon suggests that local inhibition could regulate the size of the MFR combination relayed to Purkinje neurons, without sacrificing the subsets of the combinations completely, as would occur with maximal local diversity.

### Impact of mossy fiber spacing on combinatorial diversity

Mossy fiber afferents frequently appear to terminate within clusters in the GCL. For example, both the nucleocortical pathway and the basilar pontine nuclei terminate in patches of cortex (Fig. 5A-C; Houck and Person, 2015; c.f. Huang et al., 2013). Such clustering is common, with MFs from diverse sources terminating densely along zebrin stripes, or in similarly spaced stripes within the layer (Sillitoe et al., 2010; Gebre et al., 2012; Quy et al., 2011; Gao et al., 2016; Valera et al., 2016). While MFs originating from the same nucleus do not necessarily carry the same information, single cell label supports the idea that rosettes from the same fiber terminate densely (Quy, 2011; Sultan, 2001), and physiological data support the notion that GrCs can receive information from like fibers (Jörntell and Ekerot, 2006). Before including spatial clustering in our model, we measured the patchiness of two MF pathways, analyzing the distribution of MFRs. We labeled the nucleocortical and ponto-cerebellar pathways using AAVs expressing fluorescent proteins and analyzed MFRs in Crus 1 and Lobule 6, used as representative locations (See Methods). Nearest neighbor analyses from 874 rosettes revealed that most MFRs are clustered, existing within 100 µm of another MFR from the same source (Fig. 5C).

**Figure 5.**
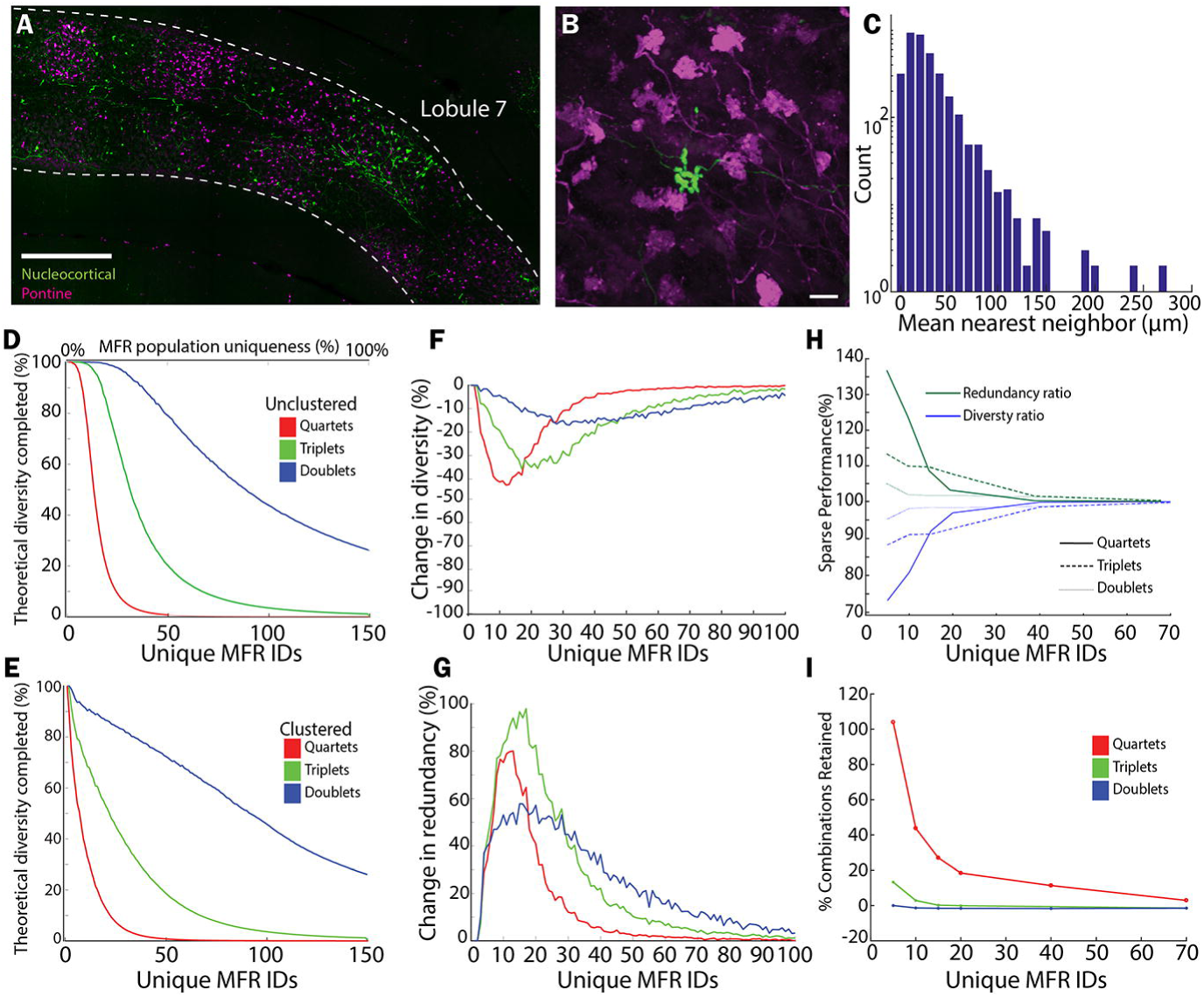
Clustered mossy fibers enhance redundancy at the expense of diversity. **A.** Representative example of patches of nucleocortical (green) and pontine (magenta) mossy fiber rosettes in mouse Lobule 7 (n = 4 mice, 8 injections). Scale bar = 200 µm. **B.** Representative example of both clustered (pontine, magenta) and sparse (nucleocortical, green) MFRs at the apex of Crus 1. **C**. Histogram of the mean distance of MFRs to their 4 closest neighbors labeled from the same injection, (n = 874 rosettes, 4 mice). **D.** Percentage of the total theoretical combinations (n choose k with replacement) produced by the modeled system, as a function of the number of different MFRs in the system. Rosettes were positioned following a uniform probability distribution, i.e no explicit rosette clustering. **E.** Percentage of maximal GrC combinatorial diversity produced as a function of MFR diversity when MFR IDs were non-uniformly represented. **F.** The difference in the diversity of MFR combinations produced with clustered vs unclustered inputs, computed by subtracting curves in Figure 5E-5D. Unclustered mossy fiber distributions support more combinatorial diversity compared to clustered mossy fiber distributions. **G.** Percent change in the redundancy of represented combinations in clustered systems vs uniform systems as a function of the number of unique MFRs in the system. At modest MFR diversity (~20 IDs), clustering nearly doubles redundancy of combinations relative to unclustered fibers. **H.** The effect of sparseness on redundancy and diversity compared to a system with n+1 uniformly distributed MFRs, plotted as a function of MFR diversity. A sparse rosette enhances redundancy but decrements diversity. **I.** Mean sparse retention index plotted as a function of MFR diversity. Sparse inputs retain more of the existing combinations that adding a uniform input.

We therefore explored the effect of these spatial characteristics on GrC combinatorial diversity in our model. We compared the GrC combinatorial diversity produced with unclustered verses clustered MFRs. To do this, we computed the number of unique GrC MFR combinations produced by the model as a function of MFR diversity relative to the number of theoretically possible combinations, based on n choose k with replacement, where k = 4, 3, or 2 for quartet, triplet and doublet combinations respectively (Fig. 5D,E, color coded by combination size). In unclustered models, MFR ID numbers were drawn at random from a uniform probability distribution. In clustered models, MFR ID numbers were drawn at random from a Gaussian-like probability distribution function, leading to over and underrepresentation of specific ID numbers in the population. Clustering MFR ID numbers attenuated the fraction of theoretical diversity produced by the model compared to unclustered inputs (Fig. 5D-F) and enhanced the redundancy of MFR combinations (Fig. 5G).

The trade-off between diversity and redundancy observed with clustering raised related questions of whether other features of mossy fiber organization affect these parameters. In addition to clustered inputs (Fig. 5A, C) we noted that, although rare, MFRs can appear hundreds of microns away from rosettes from the same source (Fig. 5B-C). We therefore asked how these sparse, numerically limited, MFRs contribute to GrC combinatorial diversity. We analyzed diversity and redundancy of GrC combinations produced in model systems mimicking these anatomical features. We modeled three systems to facilitate comparisons. The ‘baseline’ system contained MFRs with ‘n’ ID numbers uniformly distributed within the system; the ‘sparse’ system contained MFRs with ‘n’ ID numbers but additionally, a single MFR was assigned an ID number unique to the system; and the ‘expanded’ condition, in which a new ID was uniformly added to the baseline system (‘n+1’ ID numbers). Compared to the baseline system, sparse rosettes decreased redundancy and increased the diversity of combinations, similar to the effect of adding an additional input (Fig. 5H). However, compared to the expanded system, the sparse input better preserved the redundancy of combinations that were present in the baseline system, suggesting that sparse fibers increases diversity with less detriment to redundancy compared to adding another input uniformly to the space (Fig. 5I, See Methods for details).

### Impact of mossy fiber filopodia on GrC combinatorial diversity

Mossy fiber filopodia are long, thin, bouton-bearing processes that extend from MFRs (Fig. 6A-C). They have recently been shown to form synapses on GrCs (Gao et al., 2016), raising the question of their potential role in GrC combinatorial diversity. We reconstructed and analyzed pontine and nucleocortical MFRs from throughout the cerebellar cortex (Fig. 6B-D), quantifying the fraction of MFRs possessing filopodia (n = 1658 boutons), the number of boutons per filopodium (n = 84 boutons), and the distance between these boutons and the MFR (n = 84 boutons). Filopodial boutons were typically within 22 microns of the rosette, and there were between 1-4 filopodial boutons per rosette (Fig. 6D; median, 2). We found that nearly 32.2% of nucleocortical MFRs possessed filopodia, while pontine MFRs possessed filopodia at a lower rate, 11.3%, such that the likelihood of bearing filopodia for the entire population we observed was 16.3%.

**Figure 6.**
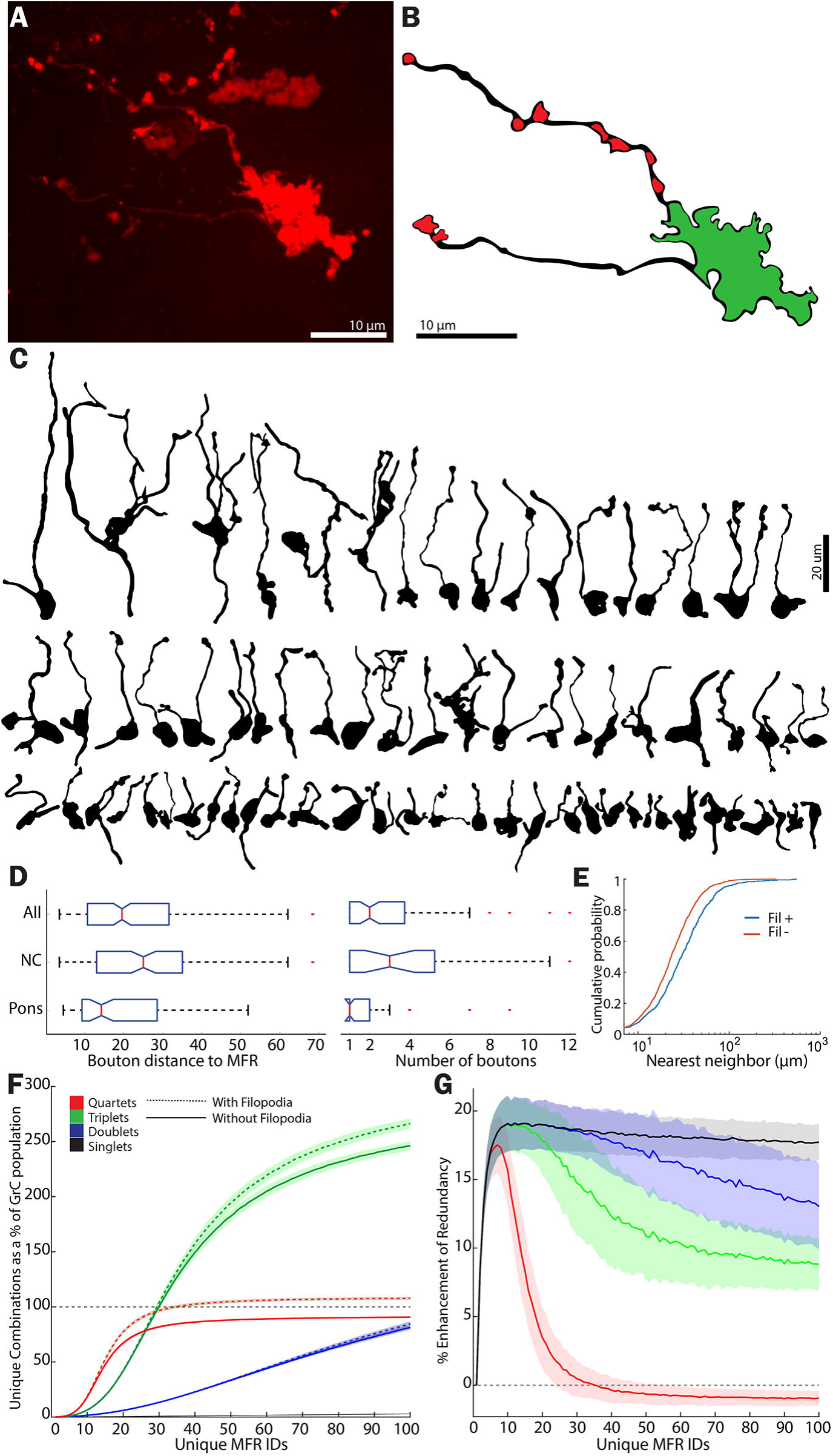
Mossy fiber filopodia enhance both combinatorial diversity and redundancy produced by GCL. **A.** A representative image of an MFR bearing filopodia. **B.** A reconstruction of MFR in *A* that illustrates the MFR in green and putative filopodial boutons in red. **C.** Reconstructions of 70 MFRs from 9 mice that bore filopodia, illustrating morphological diversity. **D.** Box plots of median and inter-quartile ranges of distances of filopodial boutons to the MFR and the number of boutons/MFR for pontine fibers, nucleocortical fibers and both sources (n = 84 MFRs, 9 mice). **E.** Cumulative distribution plot of nearest neighbors between filopodia bearing and non-filopodia bearing MFR, showing that filopodia-bearing rosettes are more sparsely positioned (n = 874 boutons, 4 mice, 8 injections). **F.** Total number of unique combinations of inputs plotted as a function of MFR diversity when 20% MFRs bore between 1-5 filopodial boutons, contacting GrCs within 22 microns in the spatially constrained model. Filopodia enhance the total number of all sizes of combinations, such that the total unique quartets outnumber GrCs. **G.** Percent increase in MFR combination redundancy seen when filopodia are added to 20% of MFRs (See methods).

We used these measurements in our GCL model, adding 1-5 synapses to GrCs located within 22 µm of a randomly selected 8-20% of MFRs, simulating filopodia. We analyzed the effect of these filopodia-like synapses on GrC combinatorial diversity and redundancy. In the model, filopodia-like extensions enhanced the diversity of GrC combination (Fig. 6E), increasing the number of unique combinations of 4 MFR inputs beyond the number of granule cells present. This effect was a consequence of increasing the number of MFRs contacting many GrCs, which was previously limited to 4 (Fig. 6F). As the diversity of MFRs increased differences between systems with and without filopodia became more pronounced (Fig 6F). MFR filopodia also enhanced redundancy of quartets in low-to-modest diversity systems, expanding the representation of individual inputs, particularly of rarer MFRs (Fig. 6G).

These modeling results indicate that filopodia mainly enhanced combinatorial diversity when MFR diversity was high, i.e. in instances when individual MFR identities are sparser. We tested whether sparseness of MFRs was related to the presence or absence of filopodia in our anatomical samples, measuring the nearest neighbors of filopodia-bearing and non-bearing MFRs. In keeping with predictions from the model, MFRs bearing filopodia were more sparsely spaced from like-neighbors. Median nearest neighbors for MFRs that did not bear rosettes was 23.1 microns, while those bearing filopodia were 30 microns apart. Differences in the distributions were highly significant (Fig. 6E; K-S goodness-of-fit test, p = 1.5 × 10^−21^). Taken together, filopodia could serve an important role in the function of the GCL to enhance representation of sparse fibers, increasing combinatorial diversity and redundancy.

## Discussion

Here we explored the effects of GCL morphological features on combination of afferents and diversification of GrCs. Our studies reveal a surprising theme of anatomical features favoring redundancy of afferent mixing rather than simply maximizing diversity. These findings raised the question of whether the spatial restrictions confer any advantage to information processing by the layer. We found that redundancy produced by spatially clustered afferents not surprisingly reduces dimensionality but enhances the capacity for temporal diversification of GrC activity. Empirical analysis of MFs validated features of the model, suggest potential evolutionary pressures structuring afferent mixing in the cerebellum.

Fundamental theoretical work on the GCL first proposed that GrCs combine afferents to support pattern separation and noise reduction by Purkinje neurons, under the guidance of climbing-fiber mediated teaching signals (Marr, 1969; Albus, 1971). Considerable empirical support for this view exists, and similar circuits are seen in diverse brain areas and species, suggesting common computational principles (Caron et al., 2014; Kennedy et al., 2014). We investigated how seemingly non-random features (i.e. patchiness) of cerebellar anatomy impact GrC combinatorial diversity. While it is perhaps obvious that every permutation of MFRs is not produced by the GCL, the specific limitations of this recoding scheme have, to our knowledge, not been described previously.

### Dendrite length and afferent diversity influence combinatorial load

We found that the short dendrites strongly limit GrC access to the full diversity of inputs to a region. Regardless of the diversity of inputs, individual GrCs access fewer than 10 different inputs (Fig. 2). The consequences of this limitation are evident when comparing the number of different MFR combinations produced by GrCs to the total number of GrCs: the diversification is submaximal, and the number of unique combinations of 4 inputs remains below the number of GrCs in the population. This indicates that as mossy fibers diversify within a region of cortex, GrCs share similar inputs, producing redundancy and reducing the dimensionality of information representation in the layer (Fig. 3; Litwin-Kumar et al., 2017).

### Benefits conferred by anatomical organization

GrC redundancy is surprising given that the computational power of combinatorial diversity is degraded with correlated and overlapping inputs (Barak et al., 2013; Rigotti et al., 2013; Litwin-Kumar et al., 2017). We speculated that over-representation of particular combinations, produced both by GrC morphology (Fig. 2) and MFR clustering (Fig. 5), may facilitate temporal expansion of GrC coding, a property hypothesized to occur in the service of learned timing (Medina et al., 2000). If similar combinations of inputs carrying specific information engage many GrCs, then inhibitory feedback mechanisms could conceivably diversify the temporal representation more effectively than if that representation is overly sparse. We tested this assumption by developing a series of equations that weighted both diversification in identity and redundancy (Fig. 3), penalizing over-representation and under-representation of combinations. We found that the spatially-organized model outperformed the random connectivity model when output featured both high dimensional recombination of inputs and temporal expandability (Fig. 3C).

These findings are interesting in light of recent studies showing denser engagement of GrCs than predicted by statistical models of the GCL (Marr, 1969). In both mice and fish, between 20 and 80% of monitored GrCs could be active within a short epoch, far greater than 1% predicted (Giovanucci et al., 2017; Wagner et al., 2017; Knogler et al., 2017). Our data suggest that this difference need not preclude combinatorial diversity as a major feature of granule layer coding but predict that this redundant code is temporally diversified, but not yet visible with relatively slow Ca^2+^ indicators used in these studies.

### Anatomical Diversity of Mossy Fibers Afferents

Although specific estimates of MF diversity are lacking, recent work employing bulk viral label of diverse precerebellar structures indicate extensive intermixing of two sources and convergence on individual GrCs (Huang et al., 2013), consistent with inferences from individual MF ramification patterns. Brainbow-labeled MFs in the cerebellar flocculus indicate highly heterogeneous fibers in a small volume (Livet et al., 2007) while rosette spacing measured in individual fibers averaged 66 ± 55 µm (Sultan, 2001). Individual fiber data therefore suggest that an average of 8 MFR duplications occur within 100 µm^3^, with an ID density of roughly 30, slightly higher than the location of peaks describing the marginal increase in GrC diversity produced by an addition MFR (between 10 and 20 IDs/247MFRs).

MF distributions from bulk labeled nuclei show much denser innervation patterns (Huang et al. 2013; Akintunde and Eisenaman, 1994; Brodal and Bjaalie, 1997; Quy et al., 2011; Shinoda et al., 1992; McCrea et al., 1977; Houck and Person, 2014; 2015). While these clusters represent ramification of different neurons, physiological data suggest that there is likely some overlap in information encoded from individual precerebellar nuclei (Jörntell and Ekerot, 2006). We explored the effect of MFR clustering on combinatorial diversity and found that redundancy is enhanced more than diversity is reduced (Fig. 5). This trade-off suggests that MF afferents may have evolved densities that exploit both combinatorial expansion by the layer and redundancy of local representation.

The nucleocortical pathway has recently regained attention as an intriguing feedback pathway within the cerebellum (Tolbert et al., 1978; Houck and Person, 2014; 2015; Gao et al., 2016). This pathway produces a fairly well-circumscribed input to the cerebellar cortex which include sparse rosettes (Fig. 5; Houck and Person, 2015; Gao et al., 2016). We found that sparse fibers increase GrC diversity, but at the cost of redundancy. However, sparse rosettes preserved more redundancy than simply adding a new MFR uniformly to the system (Fig. 5). This effect suggests that sparse inputs are able to moderately diversify systems without disrupting redundancy.

Finally, we explored the contribution of MFR filopodia to combinatorial diversity. MFR filopodia have been noted for decades (Palay and Chan-Palay, 1974; Mason and Gregory, 1984; Kalinovsky et al., 2011) and are now appreciated to form synapses on both Golgi cells (Ruediger et al. 2011) and GrCs (Gao et al, 2016). In the model, we found that filopodia allow MFRs to support more combinatorial expansion without the spatial cost of producing a new rosette. In diverse MFR systems, where individual IDs are sparser, filopodia enhanced GrC combinatorial diversity, which was in keeping with the observation that sparser MFRs were more likely to bear filopodia. Conversely, filopodia enhanced redundancy, especially in triplet and doublet systems (Fig. 6G). Our reconstructions, moreover, highlight the morphological diversity of filopodia and suggest further experiments defining the relative synaptic strength of these inputs relative to MFRs. While it is unlikely that all MFR filopodia contact GrCs, effects were graded proportional to the number of synapses added.

### Physiology of MFs and influences on combinatorial representation

Physiological data on multimodal convergence is beginning to emerge, and the details of findings in these studies highlight potential nuances to the combinatorial hypothesis. While GrCs in both mammals and fish integrate diverse information (Sawtell, 2010; Ishikawa et al., 2015; Arenz et al, 2008) the question remains whether a complete quartet of MFs is typically required for GrCs to reach threshold, or whether subsets of inputs are sufficient to drive GrC activity (Ishikawa et al., 2015; Rancz et al., 2007; Jorntell and Ekerot, 2006; Rossert et al., 2014). Recent evidence of diversity of synaptic properties from different MF types could in part explain these diverse outcomes, with some afferents having powerful driver-like effects on GrCs while other afferents are weaker but facilitate with use (Chabrol et al., 2015). This physiological diversity suggests that the number of MFs required to drive a GrC might be regulated by the specific identities of the inputs that are active.

Additionally, MFRs show LTP with theta burst stimulation (D’Angelo et al., 1999; Hansel et al., 2001), raising the question of whether increasing synaptic strength allows sub-quartet MFR combinations to drive GrCs to threshold. We found that, with increasing MFR strength, GrCs with a fixed threshold can be driven with a greater diversity of combined inputs, allowing the GCL to represent information beyond the limit imposed on quartets by the GCL population. We showed that the spatial extent of GrCs that share 3 inputs is considerably larger than those that share 4 (Fig. 4), therefore, MF-LTP could contribute both to redundant representation of information and spatially broader representations.

### Golgi cells and temporal expansion

With MFR patchiness, GrC morphology, MF physiology and MFR filopodia all enhancing GrC redundancy, the Golgi cell network becomes a critical player in regulating the size of the co-active MFR combination transmitted to Purkinje neurons, via parallel fibers activation of GoCs. Tonic inhibition via Golgi cell inhibition dynamically sets the threshold for GrCs (Brickley et al., 1996; Duguid et al., 2012; Duguid et al., 2015), and phasic inhibition produces surround inhibition and temporal sharpening (D’Angelo et al., 2013; Nieus et al., 2014; Kanichay and Silver, 2008). Because even at relatively high MFR diversity neighboring GrCs are likely to share a subset of inputs (Fig. 4), GoC inhibition is in a position to dynamically regulate the number of shared inputs driving GrCs (D’Angelo, 2008; Solinas et al., 2010).

## Conclusions

This study explored how a variety of features of granule layer organization contribute to recombining MF inputs. GCL morphology limits mixing locally, but patchy ramification patterns contribute to robust representation, which may be important for temporal diversification or sufficient representation for motor learning. Along with our findings that diversity reaches a saturation point due to anatomical constraints, our study suggests that this failure to reach maximal diversity in GCL afferents is not inherently detrimental. Finally, specializations such as sparse inputs and filopodial extensions can mitigate limitations on combinatorial diversity created by anatomical restrictions. Future studies examining the complete connectome of patches of granule layer will indicate where within the space of maximal diversity vs redundancy the cerebellar system produces, and will further illuminate the computational strategies of the cerebellum.

## Acknowledgements

Work was supported by NIH R01 NS084996-01, the Klingenstein Foundation, and the McKnight Foundation (ALP). Light microscopic experiments were performed in the University of Colorado Anschutz Medical Campus Advance Light Microscopy Core supported in part by Rocky Mountain Neurological Disorders Core Grant Number P30NS048154 and by NIH/NCRR Colorado CTSI Grant Number UL1 RR025780. We thank Dr. Brenda Houck for performing the tracer injections and histological preparation of mouse tissue for use in our modeling and analyses. We thank Drs. Joel Zylberberg and Gidon Felsen for providing helpful feedback on earlier drafts of the manuscript.

## Competing Interests

The authors declare no competing interests.

## References

Akintunde A, Eisenman LM (1994) External cuneocerebellar projection and Purkinje cell zebrin II bands: a direct comparison of parasagittal banding in the mouse cerebellum. J Chem Neuroanat. 7:75–86.

Albus JS (1971) A theory of cerebellar function. Mathematical Biosci 10: 25–61

Amaral DG, Dent JA (1981) Development of the mossy fibers of the dentate gyrus: I. A light and electron microscopic study of the mossy fibers and their expansions. J Comp Neurol. 195:51–86.

Apps R, Garwicz M (2005) Anatomical and physiological foundations of cerebellar information processing. Nat Rev Neurosci. 6:297–311. Review

Arenz A, Silver RA, Schaefer AT, Margrie TW (2008) The contribution of single synapses to sensory representation in vivo. Science. 321:977–80

Barak O, Rigotti M, Fusi S (2013) The sparseness of mixed selectivity neurons controls the generalization-discrimination trade-off. J Neurosci 33:3844–3856

Billings G, Piasini E, Lõrincz A, Nusser Z, Silver RA (2014) Network structure within the cerebellar input layer enables lossless sparse encoding. Neuron. 84:960–74.

Blomfield S, Marr D. (1970) How the cerebellum may be used. Nature. 227:1224–8.

Brickley SG, Cull-Candy SG, Farrant M. (1996) Development of a tonic form of synaptic inhibition in rat cerebellar granule cells resulting from persistent activation of GABAA receptors. J Physiol. 497:753–9

Brodal P, Bjaalie JG (1997) Salient anatomic features of the cortico-ponto-cerebellar pathway. Prog Brain Res. 114:227–49. Review.

Caron SJ, Ruta V, Abbott LF, Axel R (2013) Random convergence of olfactory inputs in the Drosophila mushroom body. Nature. 497:113–7

Chabrol FP, Arenz A, Wiechert MT, Margrie TW, DiGregorio DA (2015) Synaptic diversity enables temporal coding of coincident multisensory inputs in single neurons. Nat Neurosci. 18:718–27

Chadderton P, Margrie TW, Häusser M (2004) Integration of quanta in cerebellar granule cells during sensory processing. Nature. 428:856–6

Cheron G, Escudero M, Godaux E (1996) Discharge properties of brain stem neurons projecting to the flocculus in the alert cat. I. Medical vestibular nucleus. J Neurophysiol 76:1759–1774

D’Angelo E (2008) The critical role of Golgi cells in regulating spatio-temporal integration and plasticity at the cerebellum input stage. Front Neurosci. 2:35–46

D’Angelo E, Solinas S, Mapelli J, Gandolfi D, Mapelli L, Prestori F (2013) The cerebellar Golgi cell and spatiotemporal organization of granular layer activity. Front Neural Circuits. 7:93

D’Angelo E (2017) Challenging Marr’s theory of the cerebellum. In: Computational Theories and their Implementation in the Brain. The legacy of David Marr (Vaina LM, Passingham RE, eds.) pp 62–78. Oxford: Oxford UP

D’Angelo E, Rossi P, Armano S, Taglietti V (1999) Evidence for NMDA and mGlu receptor-dependent long-term potentiation of mossy fiber-granule cell transmission in rat cerebellum. J Neurophysiol. 81:277–87

Diwakar S, Lombardo P, Solinas S, Naldi G, D’Angelo E (2011) Local field potential modeling predicts dense activation in cerebellar granule cells clusters under LTP and LTD control. PLoS One. 6(7):e21928

Duguid I, Branco T, London M, Chadderton P, Häusser M (2012) Tonic inhibition enhances fidelity of sensory information transmission in the cerebellar cortex. J Neurosci. 32:11132–43.

Duguid I, Branco T, Chadderton P, Arlt C, Powell K, Häusser M. (2015) Control of cerebellar granule cell output by sensory-evoked Golgi cell inhibition. Proc Natl Acad Sci U S A. 112:13099–104

Eccles JC, Ito M, Szentagothai J (1967) The cerebellum as a neuronal machine. New York: Springer-Verlag

Freeman JH, Rabinak CA. (2004) Eyeblink Conditioning in Rats Using Pontine Stimulation as a Conditioned Stimulus. Integr Physiol Behav Sci. 39(3): 180–191.

Gao Z, Proietti-Onori M, Lin Z, Ten Brinke MM, Boele HJ, Potters JW, Ruigrok TJ, Hoebeek FE, De Zeeuw CI. (2016) Excitatory Cerebellar Nucleocortical Circuit Provides Internal Amplification during Associative Conditioning. Neuron. 89:645–57

Garwicz M, Jorntell H, Ekerot CF (1998) Cutaneous receptive fields and topography of mossy fibres and climbing fibres projecting to cat cerebellar C3 zone. J Physiol 512:277–293

Gebre SA, Reeber SL, Sillitoe RV. (2012) Parasagittal compartmentation of cerebellar mossy fibers as revealed by the patterned expression of vesicular glutamate transporters VGLUT1 and VGLUT2. Brain Struct Funct. 217:165–80.

Giovannuci A, Badura A, Deverett B, Najafi F, Pereira TD, Gao Z, Ozden I, Kloth AD, Pnevmatikakis E, Paninski L, De Zeeuw CI, Medina JA, Wang SS. (2017) Cerebellar granule cells acquire a widespread predictive feedback signal during motor learning. Nature Neuroscience 20:727–734

Halverson HE, Hubbard EM, Freeman JH. (2009) Stimulation of the lateral geniculate, superior colliculus, or visual cortex is sufficient for eyeblink conditioning in rats. Learn Mem. 16(5): 300–307.

Hansel C, Linden DJ, D’Angelo E. (2001) Beyond parallel fiber LTD: the diversity of synaptic and non-synaptic plasticity in the cerebellum. Nat Neurosci. 4:467–75. Review

Herculano-Houzel S. (2010) Coordinated scaling of cortical and cerebellar numbers of neurons. Front Neuroanat. 4:12.

Houck BD, Person AL. (2014) Cerebellar loops: a review of the nucleocortical pathway. Cerebellum. 13:378–85

Houck BD, Person AL. (2015) Cerebellar Premotor Output Neurons Collateralize to Innervate the Cerebellar Cortex. J Comp Neurol. 523:2254–71

Huang CC, Sugino K, Shima Y, Guo C, Bai S, Mensh BD, Nelson SB, Hantman AW (2013) Convergence of pontine and proprioceptive streams onto multimodal cerebellar granule cells. Elife; 2:e00400.

Ishikawa T, Shimuta M, Häusser M. (2015) Multimodal sensory integration in single cerebellar granule cells in vivo. Elife. Dec 29;4. pii: e12916

Jörntell H, Bengtsson F, Geborek P, Spanne A, Terekhov AV, Hayward V. (2014) Segregation of tactile input features in neurons of the cuneate nucleus. Neuron. 83:1444–52.

Jörntell H, Ekerot CF. (2006) Properties of somatosensory synaptic integration in cerebellar granule cells in vivo. J Neurosci. 26:11786–9

Kalinovsky A, Boukhtouche F, Blazeski R, Bornmann C, Suzuki N, Mason CA, Scheiffele P. (2011) Development of axon-target specificity of ponto-cerebellar afferents. PLoS Biol 9(2): e1001013.

Kanichay RT, Silver RA (2008) Synaptic and cellular properties of the feedforward inhibitory circuit within the input layer of the cerebellar cortex. J Neurosci. 28(36):8955–6

Kennedy A, Wayne G, Kaifosh P, Alviña K, Abbott LF, Sawtell NB. (2014) A temporal basis for predicting the sensory consequences of motor commands in an electric fish. Nat Neurosci. 17:416–22.

Knogler LD, Markov DA, Dragomir EI, Sith V, Portugues R. (2017) Sensorimotor Representations in Cerebellar Granule Cells in Larval Zebrafish Are Dense, Spatially Organized, and Non-temporally Patterned. Curr. Biol., 27(9):1288–1302

Kolkman KE, McElvain LE, du Lac S. (2011) Diverse precerebellar neurons share similar intrinsic excitability. J Neurosci. Nov 16;31(46):16665–74

Litwin-Kumar A, Harris KD, Axel R, Sompolinsky H, Abbott LF (2017) Optimal degrees of synaptic connectivity. Neuron 93, 1–12

Li WK, Hausknecht MJ, Stone P, Mauk MD (2013) Using a million cell simulation of the cerebellum: network scaling and task generality. Neural Netw. 47:95–102

Livet J, Weissman TA, Kang H, Draft RW, Lu J, Bennis RA, Sanes JR, Lichtman JW. (2007) Transgenic strategies for combinatorial expression of fluorescent proteins in the nervous system. Nature. 450:56–62

Mackie PD, Morley JW, Rowe MJ (1999) Signalling of static and dynamic features of muscle spindle input by external cuneate neurones in the cat. J Physiol 519:559–569.

Mapelli J, D’Angelo E. (2007) The spatial organization of long-term synaptic plasticity at the input stage of cerebellum. J Neurosci. 27:1285–96

Mason CA, Gregory E. (1984) Postnatal maturation of cerebellar mossy and climbing fibers: transient expression of dual features on single axons. J Neurosci. 4:1715–35.

Mauk MD, Donegan NH. (1997) A model of Pavlovian eyelid conditioning based on the synaptic organization of the cerebellum. Learn Mem. 4:130–58.

Marr D. (1969) A theory of cerebellar cortex. J Physiol. 202:437–70.

McCrea RA, Bishop GA, Kitai ST. (1977) Electrophysiological and horseradish peroxidase studies of precerebellar afferents to the nucleus interpositus anterior. II. Mossy fiber system. Brain Res. 122:215–28.

Medina JF, Garcia KS, Nores WL, Taylor NM, Mauk MD (2000) Timing mechanisms in the cerebellum: testing predictions of a large-scale computer simulation. J Neurosci. 20:5516–25

Nieus TR, Mapelli L, D’Angelo E. (2014) Regulation of output spike patterns by phasic inhibition in cerebellar granule cells. Front Cell Neurosci. 8:246

Palay S and Chan-Palay V (1974) Cerebellar Cortex Cytology and Organization. New York: Springer-Verlag

Palkovits M, Magyar P, Szentágothai J (1971) Quantitative histological analysis of the cerebellar cortex in the cat. II. Cell numbers and densities in the granular layer. Brain Res. 32:15–30

Quy PN, Fujita H, Sakamoto Y, Na J, Sugihara I. (2011) Projection patterns of single mossy fiber axons originating from the dorsal column nuclei mapped on the aldolase C compartments in the rat cerebellar cortex. J Comp Neurol. 519:874–99

Rancz EA, Ishikawa T, Duguid I, Chadderton P, Mahon S, Häusser M. (2007) High-fidelity transmission of sensory information by single cerebellar mossy fibre boutons. Nature. 450:1245–8

Rigotti M, Barak O, Warden MR, Wang XJ, Daw ND, Miller EK, Fusi S. (2013) The importance of mixed selectivity in complex cognitive tasks. Nature. 497:585–9

Ruediger S, Vittori C, Bednarek E, Genoud C, Strata P, Sacchetti B, Caroni P. (2011) Learning-related feedforward inhibitory connectivity growth required for memory precision. Nature. 473:514–8

Sawtell NB. (2010) Multimodal integration in granule cells as a basis for associative plasticity and sensory prediction in a cerebellum-like circuit. Neuron.66: 573–84

Schneiderman N, Gormezano I (1964) Conditioning of the nictitating membrane of the rabbit as a function of CS-US interval. J Comp Physiol Psychol 57: 188–195

Schmitz C, Hof PR (2005) Design-based stereology in neuroscience. Neuroscience 130, 813–831

Shinoda Y, Sugiuchi Y, Futami T, Izawa R. 1992 Axon collaterals of mossy fibers from the pontine nucleus in the cerebellar dentate nucleus. J Neurophysiol. 67:547–60.

Sillitoe RV, Vogel MW, Joyner AL. (2010) Engrailed Homeobox Genes Regulate Establishment of the Cerebellar Afferent Circuit Map J Neurosci 30:10015–10024

Smith MC, Coleman SR, Gormezano I. (1969) Classical conditioning of the rabbit’s nictitating membrane response at backward, simultaneous, and forward CS-US intervals. J Comp Physiol Psychol 69: 226–231

Solinas S, Nieus T, D’Angelo E. (2010) A realistic large-scale model of the cerebellum granular layer predicts circuit spatio-temporal filtering properties. Front Cell Neurosci. 4:12

Spanne A, Jörntell H. (2015) Questioning the role of sparse coding in the brain. Trends Neurosci. 2015 38:417–27.

Spanne A, Jörntell H. (2013) Processing of multi-dimensional sensorimotor information in the spinal and cerebellar neuronal circuitry: a new hypothesis. PLoS Comput Biol.;9(3):e1002979.

Steinmetz JE, Rosen DJ, Chapman PF, Lavond DG, Thompson RF (1986) Classical conditioning of the rabbit eyelid response with a mossy-fiber stimulation CS: I. Pontine nuclei and middle cerebellar peduncle stimulation. Behav Neurosci. 100:878–87

Sultan F. (2001) Distribution of mossy fibre rosettes in the cerebellum of cat and mice: evidence for a parasagittal organization at the single fibre level. Eur J Neurosci. Jun;13(11):2123–30

Tolbert DL, Bantli H, Bloedel JR. (1978) Organizational features of the cat and monkey cerebellar nucleocortical projection. J Comp Neurol. 182:39–56

Valera AM, Binda F, Pawlowski SA, Dupont JL, Casella JF, Rothstein JD, Poulain B, Isope P. (2016) Stereotyped spatial patterns of functional synaptic connectivity in the cerebellar cortex. Elife. Mar 16;5

Voogd J, Glickstein M (1998) The anatomy of the cerebellum. TINS 21, 370–375

Wagner MJ, Kim TH, Savall J, Schnitzer MJ, Luo L. (2017) Cerebellar granule cells encode the expectation of reward. Nature. 544:96–100

Welker W, Sanderson KJ, Shambes GM. (1984) Patterns of afferent projections to transitional zones in the somatic sensorimotor cerebral cortex of albino rats. Brain Res. 292:261–7

Wu HS, Sugihara I, Shinoda Y. (1999) Projection patterns of single mossy fibers originating from the lateral reticular nucleus in the rat cerebellar cortex and nuclei. J Comp Neurol. 411:97–118

